# The Object Space Task reveals a dissociation between semantic-like and episodic-like memory in a mouse model of Kleefstra Syndrome

**DOI:** 10.1101/781054

**Authors:** Evelien H.S. Schut, Alejandra Alonso, Steven Smits, Mehdi Khamassi, Anumita Samanta, Moritz Negwer, Nael Nadif Kasri, Irene Navarro Lobato, Lisa Genzel

**Affiliations:** Donders Institute for Brain Cognition and Behaviour, Radboud University, Nijmegen/Netherlands; Department of Human Genetics and Department of Cognitive Neuroscience, Radboudumc, Donders Institute for Brain, Cognition and Behaviour, 6500 HB Nijmegen, the Netherlands; Institute of Intelligent Systems and Robotics, Sorbonne Université, CNRS, Paris, France

## Abstract

Kleefstra syndrome is a disorder caused by a mutation in the *EHMT1* gene characterized in humans by general developmental delay, mild to severe intellectual disability and autism. Here, we characterized semantic- and episodic-like memory in the *Ehmt1*^*+/-*^ mouse model using the Object Space Task. We combined conventional behavioral analysis with automated analysis by deep-learning networks, a session-based computational learning model and a trial-based classifier. *Ehmt1* ^*+/-*^ mice showed more anxiety-like features and generally explored objects less, but the difference decreased over time. Interestingly, when analyzing memory-specific exploration, *Ehmt1* ^*+/-*^ show increased expression of semantic-like memory, but a deficit in episodic-like memory. A similar dissociation of semantic and episodic memory performance has been previously reported in humans with autism. Using our automatic classifier to differentiate between genotypes, we found that semantic-like memory features are better suited for classification than general exploration differences. Thus, detailed behavioral classification with the Object Space Task produced a more detailed behavioral phenotype of the *Ehmt1* ^*+/-*^ mouse model.

**One Sentence Summary:** *Ehmt1* ^*+/-*^ mice show decreased exploration and episodic-like memory but increased semantic-like memory In the Object Space Task. (143 of 150)

## Introduction

Most if not all mental disorders are accompanied by memory deficits, with the quality of the deficit depending on the overlap between the underlying circuit needed for the respected memory type and the circuit affected by the disorder. For example, semantic memories depend on cortical structures such as the medial prefrontal cortex and anterior cingulate cortex, while episodic memories are thought to rely on intact hippocampal functioning ^1-3^. Thus, larger deficits in episodic in contrast to semantic memories are expected in disorders affecting the hippocampus more than cortex. This also means that detailed characterization of the memory deficit can help predict, which circuits are affected by a disorder and lead future imaging or molecular investigations.

In animal models, most commonly simple memory paradigms are used to assess memory deficits in disease. Such tasks, e.g. contextual fear conditioning or simple object in place memory, mainly test hippocampal processing ^1^. However, just as critical for human cognition are semantic memories representing our knowledge of the world. Semantic memory is not tested by simple tasks and thus rarely assessed in animal models of disease. We recently developed a new task – the Object Space Task – that addresses this deficit: across different conditions both simple memories (episodic-like) as well as abstracted, cumulative memories (semantic-like) are tested ^4^. The task is based on a rodent’s tendency of natural exploration of new objects in an open-field environment. In the key condition of this task – overlapping – spatial configurations with two objects are presented to the animal over multiple trials per day, for 4 consecutive days. This allows the animal to accumulate information over time in order to construct an abstracted or semantic-like memory across training days. The additional advantage of this task is that it allows behavioral characterization beyond the memory measure, such as general movement patterns within an open field.

Monogenetic causes of neurodevelopmental disorders are a molecular entry point in understanding underlying mechanism and circuits of cognition. Kleefstra syndrome is a neurodevelopmental disorder characterized in humans by general developmental delay, severe intellectual disability and autism ^5-8^, caused in most cases by haploinsufficiency of the *EHMT1* gene (Euchromatic Histone Methyltransferase 1). Previous studies have shown that the heterozygous *Ehmt1* knock out mouse (*Ehmt1* ^*+/-*^) recapitulates the core features of Kleefstra syndrome.

The protein encoded by *EHMT1* (EHMT1 or GLP (G9a-like protein) acts as a histone methlystransferase, i.e. an epigenetic regulator. EHMT1 catalyzes mono- and dimethylation of histone H3 at lysine 9 (H3K9me2) ^9^, thereby working as repressor of transcription. The mouse homolog has been shown to specifically regulate the expression of several activity-dependent genes in the hippocampus following fear conditioning ^10^. Furthermore, it has been shown to be critically involved in homeostatic synaptic scaling *in vitro* and in the developing visual cortex *in vivo* ^11,12^.

*Ehmt1* ^*+/-*^ mice recapitulate core features of the human phenotype: they show delayed postnatal development and facial and cranial abnormalities that correspond to the phenotype observed in human patients ^13^. At a behavioral level, *Ehmt1* ^*+/-*^ mice display reduced exploration, increased anxiety when exposed to novel environments, and impaired social behavior ^14^. In a fear conditioning task, *Ehmt1* ^*+/-*^ mice show increased freezing and decreased extinction, another indication for increased anxiety in this mouse model ^15^. *Ehmt1* ^*+/-*^ mice perform similar to their wildtype controls in the Barnes-maze, a task that uses anxiety to the open field as a motivator, but show a deficit in simple object recognition or placement memory ^15^. Interestingly, *Ehmt1* ^*+/-*^ mice outperform wildtypes in a touch-screen pattern separation task, a process that heavily depends on the dentate gyrus in the hippocampus ^16^. Similar results on pattern separation have been demonstrated in human individuals with high-functioning autism ^17^. In contrast, Kleefstra Syndrome in humans is classically associated with severe learning disabilities. Earlier studies have shown impaired hippocampal-dependent learning in *Ehmt1* ^*+/-*^mice and impaired hippocampal physiology, including reduced excitatory connectivity between CA3 and CA1^15^. While impairments in hippocampal-dependent memory may to some extent reflect the episodic memory impairments in humans, the increased anxiety-like behavior in *Ehmt1* ^*+/-*^mice may have confounded performance in hippocampal memory paradigms in previous studies. In addition, to our knowledge, semantic-like learning abilities in these animals have not yet been tested in those mice.

Thus, to further characterize both semantic-like and episodic-like memory processes in *Ehmt1* ^*+/-*^ mice, we assessed performance of these animals in the Object Space Task^4^. The task contains both an episodic-like as well as semantic-like memory condition. In addition to the conventional behavioral analysis, we modeled the learning behavior over training days as well as build a classifier for individual trial behaviors automatically extracted from the videos with in-house deep learning networks based on DeepLabCut ^18^. *Ehmt1* ^*+/-*^ mice showed overall decreased exploration of the objects and more corner sitting but expressed increased semantic-like memory compared to controls. In contrast, *Ehmt1* ^*+/-*^ mice showed reduced episodic-like memory. Computational modelling revealed that the difference in semantic-like memory stems from a change in memory expression and not learning rate. Finally, using the classifier we showed that behavioural measures from video analysis of individual trials allow the automatic identification of genotype. The classifier performed best in our semantic-like memory condition using memory-specific features (discrimination index). Thus we could show that our Object Space Task allows for the in-depth characterization of innate behavior as well as episodic- and semantic-like memories in animal models of disease that can guide future circuit investigations.

## Results

To assess semantic-like and episodic-like memory in the *Ehmt1* ^*+/-*^ mouse model, we used the Object Space Task ^4^. In this task, mice are allowed to explore sets of objects (two identical objects, each trial a different set) in an open-field box (75cmX75cm) that are placed according to one of three conditions (see Fig. 1): In the random condition – the negative control – the objects are in semi-random configurations (randomness restricted by counterbalancing). In the stable condition the two objects always remain in the same positions until the final trial (trial 21, test) when one of the objects is moved. The stable condition tests memory of the most recent event, i.e. episodic-like memory. Finally, the key condition, overlapping, always has one object at the same position, while the other object is placed in one of the three other corners (stable location is counterbalanced across animals). Thus over time and trials, animals build up a cumulative or semantic-like memory resulting in a preference for the less often shown location ^4^. At the test trial, the same configuration as the final training trial is used, thus controlling for episodic-like memory. In this study, mice (WT n=31, *Ehmt1* ^*+/-*^ n=24, 8 weeks, male) were first habituated (1 week) and then trained (3 weeks, each week 1 session of 20 training and 1 test trials, 5 trials a day) in the task (Fig. 1). During the training period the animals went through the 3 conditions in a counterbalanced order.

**Fig. 1.**
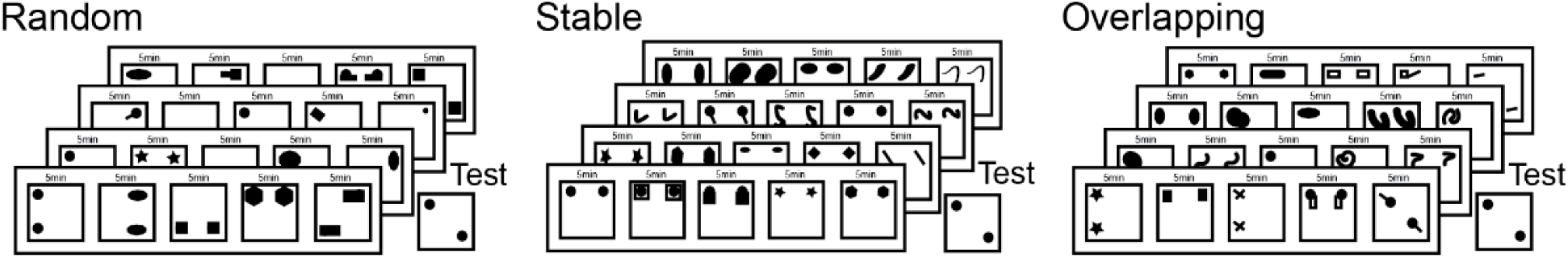
Fig. 1. Study Design. **Study Design:** Across three weeks animals are trained in three conditions in a counterbalanced order with Mo-Thu training 5 trials/day (each 5min) and Fri test (1 trial 10min). The three conditions differ in the underlying statistical regularities of object placement. Random: semi-random (constricted to equal number of occurance across the week) placement; Stable: during training the same two locations, one object moved at test; Overlapping: one location always contains an object, the second object is in one of the other three corners. Final training trial and test trial have the same configuration to control for episodic-like memory. The stable and moved location identities were counterbalanced across animals to control for general location preferences.

### General Exploration Differences: Decreased exploration in Ehmt1 ^+/-^mice

Initially, we focussed our analysis on differences in explorative behaviour and compared total exploration time, total count of object visits, and average exploration bout length (scored manually, blinded to condition and genotype). *Ehmt1* ^*+/-*^ mice show decreased total exploration time (p=0.004), however this difference decreased over time (each session is one week, session X gene interaction p=0.001, Fig. 2). The difference in overall exploration time is explained by a lower number of visits to each object (p=0.003), while average length of each individual exploration bout did not differ (p=0.223). Interestingly, over time (both within a week as well as across weeks) bout length increased independent of genotype (p=0.027), while the number of visits decreased over time in WT. Thus the convergence of total exploration time over sessions is explained by sustained number of visits with increasing bout length in *Ehmt1* ^*+/-*^and decreased number of visits with increasing bout length in WT. Total exploration time, count of explorations and bout length did not differ significantly across training conditions for either genotype.

**Fig. 2.**
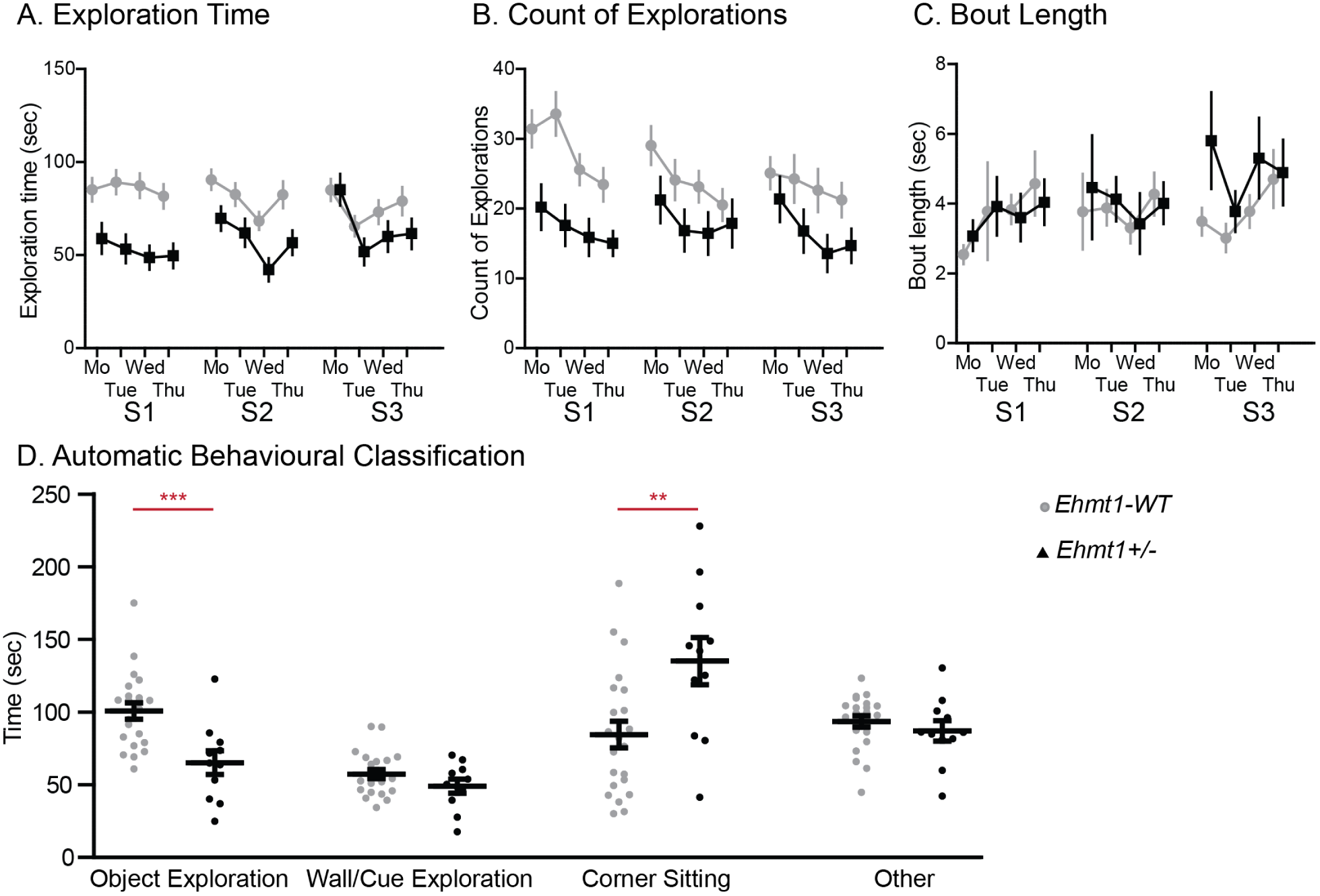
Fig. 2 General Exploration for WTIE*hmt1*+/-. **General Exploration for WT/ *Ehmt1* ^*+/-*^ Shown is A**. exploration time **B.** count of exploration bouts and **C.** average bout length across the three weeks/sessions (S1, S3, S3) of training. Each day averaged across 5 trials. *Ehmt1*^*+/-*^ show decreased exploration time due to decreased number of object visits, but this difference decreases over the three week training period. **Exploration Time** sessionXgene F_2,104_=7.3 p=0.001 (linear constrast p=0.002), day F_3.2,208_=17.3 p<0.001 (linear contrast p<0.001), sessionXday F_5.5,416_=4.6 p=0.001, gene F_1,52_=9.0 p=0.004; **Count** session F_2,82_=8.2 p=0.001 (linear contrast p=0.001), sessionXgene F_2,82_=5.6 p=0.005 (linear contrast p=0.019), day F_2.5,104_=26.1 p<0.001 (linear contrast p<0.001), dayXgene F_3,123_=4.0 p=0.009 (quadratic contrast p=0.026), sessionXday F_4.5,185_=2.2 p=0.048, sessionXdayXgene F_6,246_=2.5 p=0.022, gene F_1,41_=9.7 p=0.003; **Bout Length** session F_2,82_=3.8 p=0.027 (linear contrast p=0.026), day F_2.5,123_=3.4 p=0.025 (linear contrast p=0.021), dayXgene F_3,123_=3.0 p=0.031 (linear contrast p=0.008), sessionXday F_4.5,183_=4.0 p=0.003, gene F_1,41_=1.5 p=0.223. **D**. shows the different behaviours extracted from the automatic classification of a sub-set of videos. *Ehmt1* ^*+/-*^ spend less time exploring objects but more time sitting in the corners Behavior F_3,93_=13.9 p<0.001, BehaviorXgene F_3,93_=8.3 p<0.001, object exploration p=0.001, corner sitting p=0.006). Data shown as mean and SEM.

To further asses what the animals were doing when they were not exploring objects, we extracted different behaviors in an automatic way (object exploration, wall/cue exploration, corner sitting) from a randomly selected sub-set of the video data (1764 of 3465 trials) with in-house deep-learning networks based on DeepLabCut ^18^.

The first, in-house model used Kinetic Action Recognition to extrapolate when the mice explored objects. The model was trained on in-lab available action labeled Object Exploration video data. The second model used pose estimation from DeepLabCut to extract the actions wall/cue exploration and corner sitting (because no labeled data are available for these actions). In the end, the predicted actions of both models are frame-wise concatenated for each video trial. The code implementation for this section can be found at https://github.com/Iglohut/autoscore_3d. Automatic behavioral classification confirmed the difference in object exploration time (p=0.001) and revealed that *Ehmt1* ^*+/-*^animals spent more time sitting in corners (p= 0.006, Fig. 2D).

In sum, the initial difference in exploration time due to decreased object visits and more corner sitting in *Ehmt1*^*+/-*^ *mice*, could indicate increased anxiety-like features (reluctance to leave corner for an object visit). This decrease in exploration and increase in sitting has been shown before ^14^, but we could add that the anxiety decreases with time and habituation over the weeks of training.

### Memory Specific Differences: Decreased Episodic-like and Increased Semantic-like Memory in Ehmt1^+/-^ mice

After characterizing general exploration features, we assessed memory performance by calculating the discrimination index for each exploration time, count of explorations and bout length (DI={moved-stable}/{moved+stable}). In our episodic-like memory condition (stable) only WT were above chance at test (in the discrimination indicies for exploration time, count, bout length) and not *Ehmt1*^*+/-*^ mice (Fig. 3A). As expected in our negative control condition (random) neither genotype was above chance at test (Fig. 3B).

**Fig. 3.**
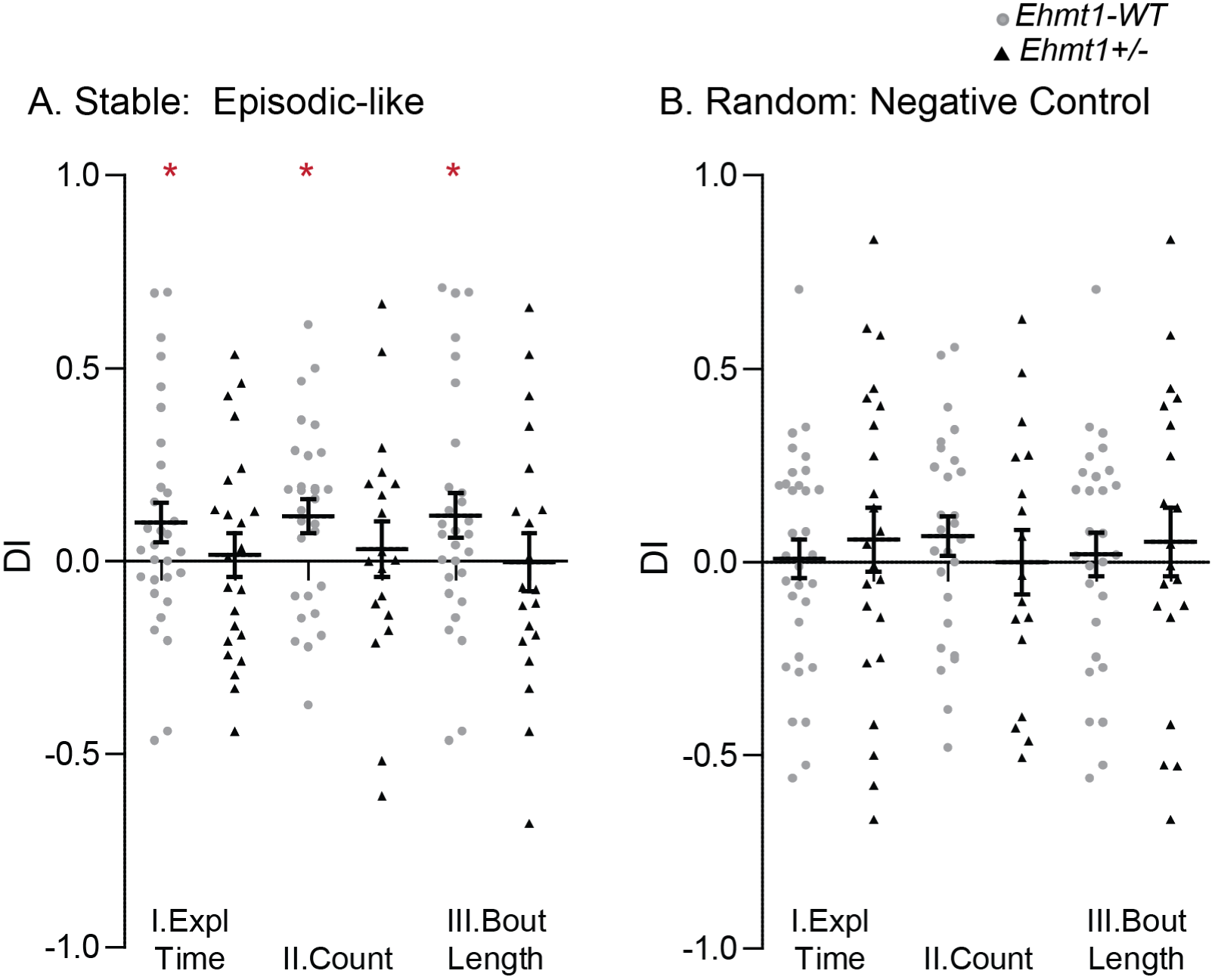
Fig.3 Episodic-Like Memory for WT/*Ehmt1*+/- (Test) **Episodic-like Memory (stable and random conditions):** Shown are the discrimination indicies at test (trial 21) calculated for I. exploration time II. count of exploration bouts and II. average bout length for both stable and random. Only WT and not *Ehmt1* ^*+/-*^performed above chance level in our episodic-like memory test (stable). T-**Test to chance in stable:** *Exploration Time* WT wilcoxon rank test p=0.03; *Count* WT t-test t_28_=2.6 p=0.01; *Bout Length* WT t-test t_28_=2.1 p=0.05. Data shown as mean and SEM.

In our semantic-like memory condition – overlapping – we would expect a positive discrimination index during training (trials 1-20) as well as test. In contrast to stable with static objects and random, there is an aligned pattern across animals in overlapping already during training and the animals develop a preference for the less-stable location (resulting in a positive DI, Fig. 4 I). At training both WT and *Ehmt1* ^*+/-*^ showed a discrimination index above chance expressed across all three variables (exploration time, count, bout length) but *Ehmt1*^*+/-*^ showed slightly higher discrimination index values especially during training (only significant p<0.05 for exploration time) indicating increased memory expression (Fig. 4). At test both WT and *Ehmt1*^*+/-*^ showed above-chance discrimination index in the overlapping condition, but due to high variability only significantly so (p=0.027) in the count of explorations measure (Fig. 4 III). Thus both WT and *Ehmt1*^*+/-*^ have semantic-like memory expression, with *Ehmt1*^*+/-*^ showing a slightly stronger effect (overlapping condition). In contrast, only WT express episodic-like memory at test (stable condition). This decrease in episodic-like memory in stable replicates findings in a one-trial object recognition and location paradigm in these mice ^15^.

**Fig. 4.**
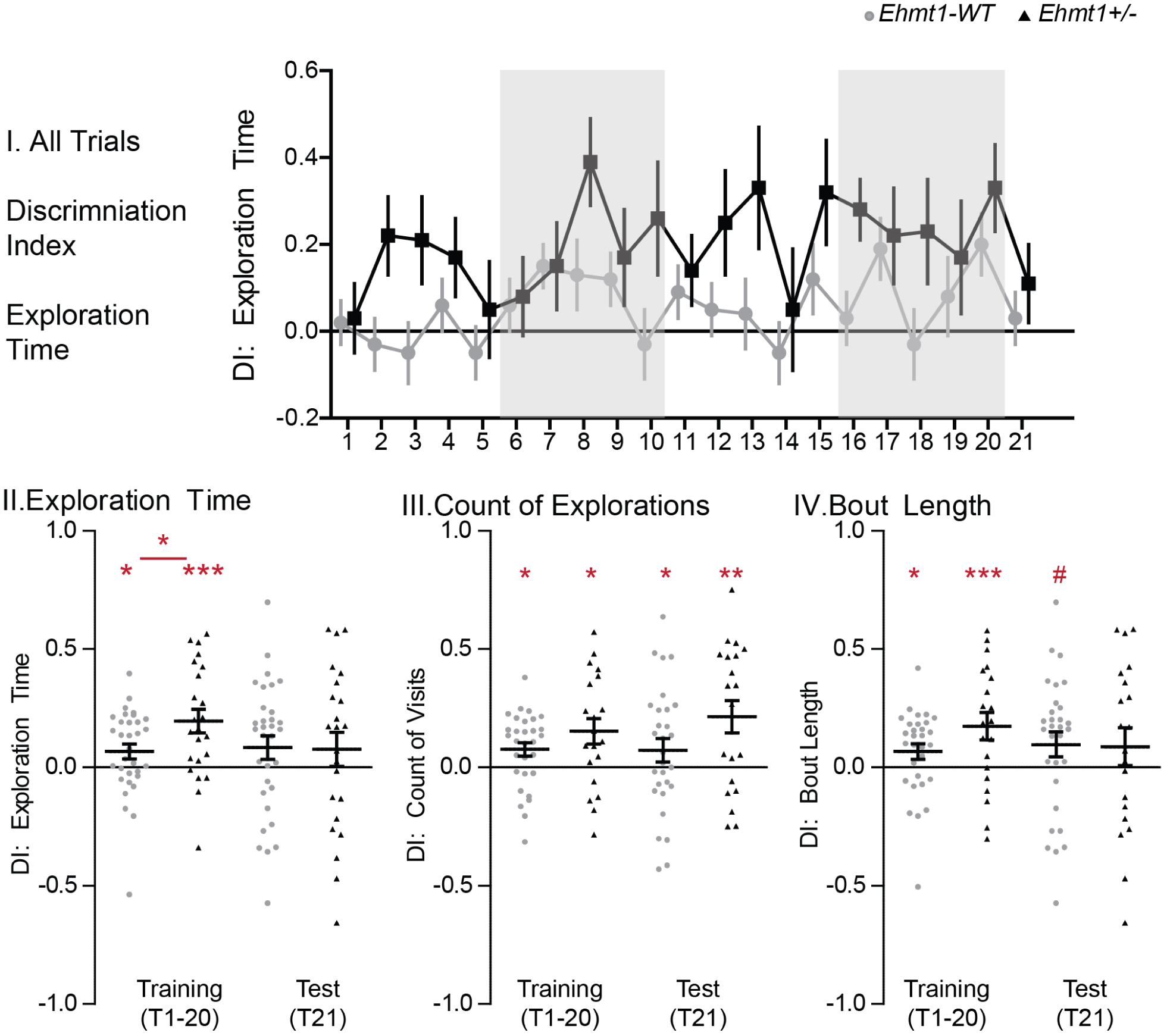
Fig.4 Semantic-Like Memory for WT/*Ehmt*+/- (Overlapping) **Semantic-like Memory (Overlapping condition):** Discrimination Index for WT/ *Ehmt1* ^*+/-*^. Shown are the discrimination indecies for training (trials 1-20) and test (trial 21) calculated on I. exploration time for each trial seperatly, and averaged across training II. exploration time, III. count of exploration bouts, and IV. average bout length. **T-test** *Ehmt1* ^*+/-*^**vs WT** overlapping pt t_53_=2.3 p=0.027; **Test to chance Training:** *Exploration Time* WT wilcoxon rank test p=0.02; *Ehmt1* ^*+/-*^t-test t_22_=4.0 p=0.0007; *Count* WT t-test t_26_=2.7 p=0.01; *Ehmt1* ^*+/-*^t-test t_19_=2.8 p=0.01; *Bout Length* WT wilcoxon rank test p=0.015; *Ehmt1* ^*+/-*^t-test t_19_=3.0 p=0.008; **Test to chance Test:** *Count* WT t-test t_25_=2.3 p=0.03; *Ehmt1* ^*+/-*^t-test t_19_=3.1 p=0.006; *Bout Length* overlapping WT t-test t_26_=1.9 p=0.066; Data shown as mean and SEM, grey shading in I for individual training days.

### Modelling learning: Ehmt1 ^+/-^ mice show same learning rate as WT but increased semantic memory expression

The decreased overall exploration seen in the *Ehmt1* ^*+/-*^ mice could have confounded the difference seen in the discrimination index in the overlapping condition. Thus, to further disentangle these effects and to characterize the build-up of a memory trace and its expression, we applied a computational model ^4^ that progressively learns place-object associations and makes decisions about which proportion of time to spend exploring each object in order to minimize uncertainty about these place-object associations. The model employs 2 parameters: a learning rate *α* which determines the balance between recent and remote memories; an inverse temperature *β*, which determines the balance between neophilic (preference for more novel/uncertain object location, positive values) and neophobic (aversion for more novel/uncertain object location, negative values) exploratory behaviors, and thus measures memory expression. Values around 0 indicate that the behaviour is not driven by memory. We fitted the model separately for each animal and each condition (stable, random, overlapping), in order to observe potential differences in the optimized parameters between conditions. Thus together *α* and *β* let us disentangle if animals actually have a better memory and/or learn faster (*α*) or just express their memory more independent of memory strength (*β*).

As with the conventional discrimination index, *β* values took more extreme either positive of negative values in *Ehmt1* ^*+/-*^ animals, which is why we took the absolute value of *β* as the key parameter for the following analyses: we want to characterize the strength of memory expression independent of neophobic or neophilic tendencies. Interestingly, *Ehmt1* ^*+/-*^animals showed significantly higher absolute *β* values but only in the overlapping condition (Kruskal-Wallis, Chi2 = 4.09, p = 0.0432, Fig. 5A). In contrast, learning rate *α* did not differ between genotype for any condition (Fig. 5B). We did replicate the finding from ^4^ that overall, the stable condition shows higher learning rates (more recent memory thus more episodic-like) in contrast to the random and overlapping conditions, which show smaller *α*’s (more influenced by remote memory thus more semantic-like).

**Fig. 5.**
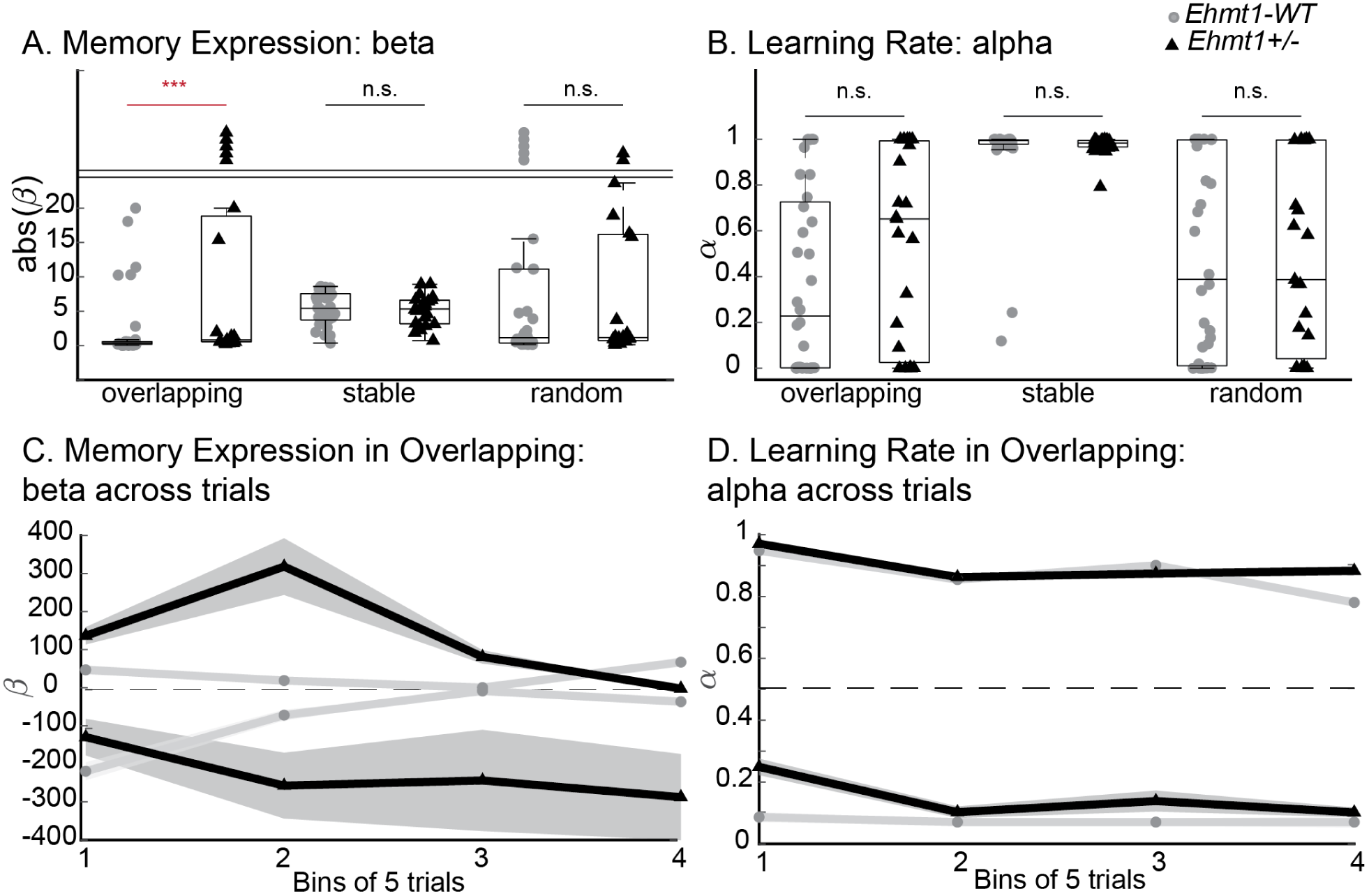
Fig. 5 Modelling Memory Expression and Learning Rate for WT*/Ehmt*+/-. **Modelling Memory Expression and Learning Rate:** A. Absolute memory expression (*β*) only showed a significant difference between the two genotypes in the overlapping condition (Kruskal-Wallis, Chi2 = 4.0885, p = 0.0432, values above 25 stacked above the line). B. No differences between WT and *Ehmt1* ^*+/-*^was seen in the learning rate (*α*). Focussing on the overlapping condition memory, memory expression (C) and learning rate (D) plotted by bins of 5 trials with groups split by genotype and by neophilic/phobic preference (pos/neg *β*) or by *α*=0.5. WT grey, *Ehmt1* ^*+/-*^black.

Next, we reran the model only within a day (bins of 5 trials) to characterize the development over the week of training. Memory expression (absolute *β*) decreased over the week (Fig.5C), which may explain the decreased memory expression at test seen in the discrimination index of total exploration time (Fig. 4). However, the learning rate remained constant (Fig. 5D).

To conclude, our model shows that WT and *Ehmt1* ^*+/-*^ learned at a similar rate. Consequentially, differences in discrimination index are based on increased memory expression in *Ehmt1* ^*+/-*^. However, this difference is specific to semantic-like memories that show underlying statistical regularities as tested in our overlapping condition.

### Automatic Behavioural Scoring and Classifier for WT/ Ehmt1 ^+/-^

On a per-session level including all 21 trials, we have shown that Ehmt1^+/-^ mice explore objects less and show increased memory expression, which is specific to our semantic-like memory condition. Next, we analyzed discrete behaviors on a trial-by-trial level: We trained two classifiers on multiple behavioral features extracted from the video data automatically with deep-learning methods as explained in the Materials and Methods section ^18^. We included both general exploration features (e.g. min explore time) as well as memory specific features (e.g. discrimination index, total 45 features, see Tab. 1).

**Table 1.** Model Features: Features extracted from the sequence data per trial. In total 45 features were calculated (above features for each object (i) and min (n) respectivly thus 13 feature description result 45 specific features).

| *Feature* | *Description* |
| --- | --- |
| First object | Whether the first object explored was either object 1 or object 2 |
| First object latency | Latency to first exploration of any object |
| Stay<sub>i</sub> | Likelihood of exploring object i next, after having last explored object i. Where i is either 1 or 2. |
| SteadyState1(SS<sub>1</sub>) | The probability of exploring object 1 after many explorations |
| Perseverance Index | Index ranging from -1 to 1 represent the tendency of switching between objects during exploration as -1, the tendency to reexplore the same object during exploration as 1, and no tendency as 0 |
| n_transitions | Total number of transitions made between objects during the whole trial. Per minute i, the total number of explorations of any object up until that minute |
| mini_n_explore | Per minute i, the total number of explorations of object j up until that minute |
| mini_n_explore_object |  |
| j mini_time | Per minute i, the total time any object was explored up until that minute |
| mini_object<sub>j</sub>_time | Per minute i and object j, the total time that object was explored up until that minute |
| bout_time | Mean time of explorations of any object |
| bout_time_obj<sub>j</sub> | Mean time of explorations of object j |
| mini_DI | Per minute i, the discrimination index (DI) up until that time representing a preference for exploring object 2 as -1, a preference for exploring object 1 as 1, and no preference as 0 |

Extracted features were fed into two classifiers: Random Forest and XGBoost. To test whether the AUROC (Area Under the Receiver Operating Curve, ROC) performance metric of each model was above the expected value of chance, both models were tested under the permutation distribution (see Materials and Methods section). Performance of each model for each condition (all pooled, overlapping, stable, random) are depicted in Fig. 6A. All models except the ones in the random condition have an AUROC > 0.61 that is significant (∀i: pi < 0.001) under the permutation distribution. With the classifier performing at a similar level when only including the data of the overlapping condition in itself as using all data from all conditions, despite the data-set being only 1/3 of the size of all pooled.

**Fig. 6.**
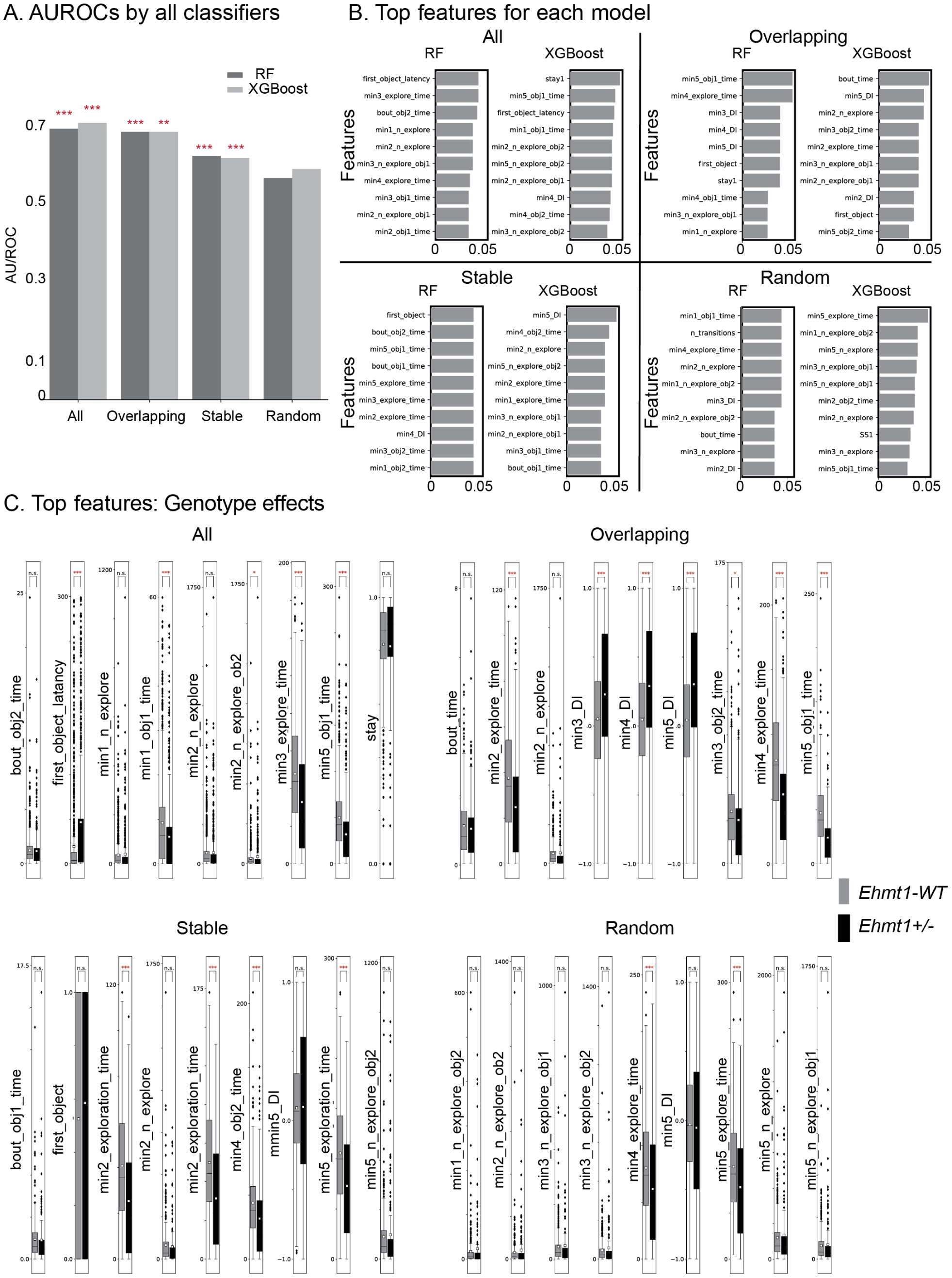
Fig. 6: Automatic Behavioural Classifier for WT/*Ehmt1*+/-. **Classifier for WT/ *Ehmt1* ^*+/-*^: A**. AUROCs by all classifiers for all subsets of data (all conditions, stable, overlapping, random). Asterisks represent the p-value of its respective AUROC under the permutation distribution: **B**. Top 10 features and their relative feature importance per model for all subsets of data (all conditions, stable, overlapping, random). **C**. Box plots of the top 9 features per model for all subsets of data (all conditions, stable, overlapping, random). The white square represents the mean. The difference of the mean between genotype was tested for each feature under the permutation distribution. * = (p < .05), ** = (p < .01), *** = (p < .001).

Furthermore, inspection of each significant model’s ROC-curve shows that all models have relatively high specificity relative to their sensitivity (sensitivity≈ 0.35 - 0.49, specificity≈ 0.87-0.89). For the models trained on trials in the random condition, neither model (Random Forest, AUROC ≈ 0.56, p > 0.05 XGBoost, AUROC ≈ 0.59, p > 0.05) could predict the genotype based on single trials.

Top ten features’ relative importances (Fig. 6B) were tested under the permutation distribution and also compared between genotype (Fig. 6C). A high relative feature importance means that a model uses that specific feature and all its potential interactions with other features. Interestingly, top features used in all, stable and random conditions were based on general exploration features such as total number of exploration bouts in the first 2min of the trial. In contrast, such general exploration features became less weighted in the overlapping condition, in which memory-based features gained importance (e.g. DI at 3 min/4 min/5 min).

The increased weights of memory-based features would further support that while we see genotype-specific differences in general exploration features, differences in semantic-like memory expression are more prominent and thus here drive automatic classification of WT and *Ehmt1* ^*+/-*^.

## Discussion

In this study, we used the Object Space Task to characterize the behavioral phenotype of the *Ehmt1* ^*+/-*^ mouse model in detail. The task tests general as well as memory-specific exploration features, and contained both an episodic- and semantic-like memory condition. By combining conventional behavioral analysis with automatic behavioral analysis via deep-learning networks, computational modelling of learning behavior across days, and a trial-by-trial behavioral classifier, we could elucidate a variety of behaviors influenced by genotype and their relative importance.

*Ehmt1* ^*+/-*^animals showed 1) decreased total exploration time due to decreased number of object visits and increased corner sitting. Over training sessions, both parameters approached WT levels. 2) In contrast to WT, *Ehmt1*^*+/-*^ show no episodic-like memory, but 3) increased semantic-like memory expression. 4) Modelling revealed that *Ehmt1* ^*+/-*^and WT showed similar learning rates in all conditions, but *Ehmt1* ^*+/-*^ showed increased memory expression (*β*) in comparison to WT. This difference was specific to the semantic-like memory condition (overlapping). 5) Likewise, computational video analysis with two different classifiers could differentiate between *Ehmt1* ^*+/-*^and WT on a trial-by-trial basis. Relative weights revealed that memory-features in the semantic-like condition showed a larger difference and drove the classifier more than general exploration features.

### General Exploration Differences

Initially *Ehmt1* ^*+/-*^ mice visited the objects less frequently and remained more in the corners than WT, perhaps indicating increased anxiety. Balemans et al also showed that *Ehmt1* ^*+/-*^ mice display impaired social behavior as well as reduced exploration and increased anxiety when exposed to novel environments ^14^. However, the Object Space Task allowed us to add to this finding, in that this increased anxiety seems to habituate over time: in the third week of training (fourth week of box exposure) there was no difference between WT and *Ehmt1* ^*+/-*^. Thus, *Ehmt1* ^*+/-*^ mice do not show persistently increased anxiety, instead they only need more time to habituate to novel situations.

The difference in exploration behavior is sufficient for a classifier to determine genotype on a trial-by-trial basis. Especially the amount of exploration by minute 2 seems to drive the classifier when considering the stable or random condition. In contrast, when determining genotype in the overlapping condition, memory-related behaviors (discrimination index at different time points) outweigh general exploration features. This indicates that while the genotypes differ in general exploration behaviors, they differ even more in semantic-like behavior.

Automatic classification of genotype based on video analysis is becoming more popular and shows great potential for monitoring treatment outcomes in pre-clinical studies. Both 3D and 2D video techniques can be used ^18,19^. Our findings highlight the importance of considering which behaviors are recorded with the video-data: Behaviors with higher cognitive demands (such as our overlapping condition) may be more sensitive to genotype differences as we can show here.

### Memory-specific Exploration Differences

In contrast to WT, *Ehmt1* ^*+/-*^ mice did not perform significantly above chance at test in our episodic-like memory task (stable). This effect is similar to previous reports ^15^, in which *Ehmt1* ^*+/-*^ mice demonstrated significantly reduced discrimination index compared to wildtype controls in a one-event object location memory test. This memory deficit may be due to hippocampal dysfunction since episodic-like memory relies on this circuit ^1^. Other differences in hippocampal function have been found between *Ehmt1* ^*+/-*^ mice and littermate controls, such as increased excitability in CA1 neurons ^15,20^.

Interestingly, *Ehmt1* ^*+/-*^ expressed a stronger semantic-like memory than wildtype controls. Initial evidence for this effect came from the conventional behavior analysis and we confirmed by modelling learning rate and memory expression. The learning model also revealed that especially absolute *β* (memory expression) differed between genotypes and thus the strength of memory expression independent of neophobic and neophilic tendencies. This also explains why conventional analyses only weakly showed this difference in memory expression: *Ehmt1* ^*+/-*^ mice show more extreme discrimination values (closer to −1 and 1) than the WT. With the mix of neophobic and neophilic tendencies that is present in both genotypes, the average discrimination index is similar between genotypes, disguising the difference in memory expression. In short, a distribution from −1 to 1 in *Ehmt1* ^*+/-*^ (and −0.5 to 0.5 in WT) will result in overall the same average but difference in standard deviation.

In contrast to memory expression, learning rate did not differ between the genotypes. Thus *Ehmt1* ^*+/-*^did not simply show better memory or learned faster, instead the same memory strength was just expresed behaviourally more in these animals. Elucidating this difference is another argument why it is important to go beyond conventional analysis of behaviour when performing phenotyping.

Notably, other experiments also showed comparable performance between *Ehmt1*^+/-^ and WT in some behavioral tasks that typically involve the cortex as well as the hippocampus, including spatial learning in the Barnes Maze ^15^. Additionally, pattern separation learning in a touch-screen task is superior in *Ehmt1* ^*+/-*^ mice ^16^. It is highly likely that successful extraction of statistical regularities in the overlapping condition of the Object Space task requires both the hippocampus and the neocortex ^3,21,22^

In sum, the dissociation between a deficit in episodic-like and intact semantic-like memory, may hint at a selective hippocampal and not cortical dysfunction in the *Ehmt1*^*+/-*^ mouse model. Future experiments combining the Object Space Task with neural recordings may help elucidate the underlying mechanism.

### Implications for Phenotype

Autism is a complex condition characterized by impaired social behavior, perseverant behaviors and communication deficits^23^. Memory processes in autism are affected as well. Whereas episodic memory deficits have been found consistently ^25,26^, semantic memory abilities and gist extraction may be equal or even superior in individuals with autism in comparison to control subjects ^27-30^. This dissociation of deficit across memory type, is reminiscent of our findings in the *Ehmt1*^*+/-*^ mouse model, where we also found a deficit in episodic-like but intact semantic-like memory. Autistic children are known for their increased need to pattern-separate: toys will often be arranged by color and size. Perhaps the increased semantic-like memory expression seen in *Ehmt1*^*+/-*^mice, which has underlying statistical regularities, is a reflection of this characteristic in mice. Increased anxiety especially in novel situations, is often seen in autism as well. In Ehmt1 mice, this phenotype was expressed in the decreased number of visits to the objects and more corner sitting in the task. The effect alleviated over time, indicative of habituation.

Overall, the more detailed characterization of memory and behavior with the Object Space Task in this model provides initial evidence that the phenotype may be reflecting autism more than intellectual disability as no deficits in semantic memory were seen. This is in contrast to the human, in which intellectual disability features are more prominent but autism features are also present ^24^. To further classify the autistic features in the mouse model, tasks testing complex social interactions should be employed next.

### The Object Space Task for Phenotyping

In this study, we used the Object Space Task for detailed behavioral characterization of the *Ehmt1*^*+/-*^ mouse model. The advantage of the task in comparison to other behavioral assays is that two types of memories are tested in a controlled and comparable setting (episodic-like and semantic-like), that due to differences in underlying circuity can show a dissociation of deficits in neurological disorders. In addition to memory specific effects, general exploration, and movement patterns in an open-field environment can also be characterized in this task. When evaluating a behavioural phenotype, it is critical to avoid confounding effects such as increased anxiety or decreased mental flexibility. Thus, one should be cautious with one-trial evaluations of behavior. These factors are controlled for in the Object Space Task and thus more nuanced phenotypes that normally would be occluded by confounding factors can be measured. Finally, we could also show that the task can easily be combined with automatic video analysis, modelling learning behavior as well as a trial-by-trial classifier that allow the in-depth characterization of phenotype beyond conventional behavioral measures.

### Conclusion

In sum, we could show that Ehmt1^+/-^ mice show increased semantic-memory compared to controls, but show deficits in episodic-like memory and increased habituation time to environments. We did so by combining conventional behavioral analysis with a session-based learning computational model and a trial-based classifier in the Object Space Task.

## Acknowledgements

We would like to thank the bachelor and master students, who ran the behavioural experiments: Joanne Igoli, Gülberk Bayraktar, Nikkie Cornelisse, Luc Nijssen, Elif Gizem Kain, Betul Tamer, Stefanos Louizo, Joris van Hout, Joost Pleune, Anne Mulder, Alysha Maurmair, Niamh Hemingway. Ronny Eichler for technical support and Veronica Pellis for labelling of videos for DeepLabCut. This work was supported by the Branco Weiss Society in Science grant to Lisa Genzel.

## Author contributions

E.S. helped design and perform the experiments, supervised the students and did the conventional behavioral analysis; AA, AS, INL helped execute the experiments and supervise the students; S.S. made the classifier, M.K. made the learning model, M.N. performed the genotyping, N.N.K. provided the mouse mode,; L.G. designed the experiment, supervised the students, performed the conventional analysis, supervised the model and classifier, and wrote the first draft of the manuscript. All authors helped revise the manuscript.

## Materials and Methods

### Subjects

Male wildtype and Ehmt1^+/-^ mice (littermates, bred in-house), 12-16 weeks of age at the start of behavioral training were group housed with ad libitum access to food and water. Mice were maintained on a 12hr/12hr light/dark cycle and tested during the light period. In compliance with Dutch law and Institutional regulations, all animal procedures were approved by the Centrale Commissie Dierproeven (CCD) and conducted in accordance with the Experiments on Animal Act.

### Behavioral training

Habituation and behavioral training has been described previously in ^4^. Briefly, all animals were extensively handled before the start of habituation. Mice were habituated to a square arena (75cmx75cm) for 5 sessions within 5 days. In the first session, mice were placed in the arena together with all cage mates for 30 minutes. In the remaining sessions, mice were placed in the arena individually for 10 minutes. In the final two sessions, two objects (made from Duplo blocks, not used in the main experiment) were placed in the arena and animals were allowed to explore.

Animals were trained on all three conditions: stable, overlapping and random. Conditions and locations were counterbalanced among animals and sessions, and the experimenter was blinded to the condition and genotype. At the beginning of each 5-day session, cues were placed on the walls inside the box and at least one 3D cue was placed above one of the other walls. Cue distribution was intentionally non-symmetric. A camera was placed above the box to record every trial and to allow for online scoring of exploration time with our in-house scoring program, the Object Scorer. In each condition, animals were allowed to explore two objects for 5 minutes with an inter-trial interval of 30min. Before the beginning of each sample trial, the box and the objects were thoroughly cleaned with 70% ethanol. Each sample trial consisted of a different pair of matching objects varying in height width, texture and material. Objects were never repeated during the training period of one condition (1 session). The test trial, 24hrs after the last sample trial, consisted of again two objects and animals were allowed to explore for 10minutes, however only the initial 5min were used for analysis. The Object Scorer software (described in Chapter 2) was used for online scoring and extraction of exploration times during all trials.

For the overlapping condition session, one object location was designated as the ‘stable’ object location, indicating that in each trial over the course of the entire session one object was positioned in this location. The other object location (‘less stable’ or ‘moved’ object location), was positioned in any of the other possible object locations, in a pseudo-random fashion. In the stable condition, two objects remained in the same location across all sample trials but one object moved during test. Finally, in the random condition objects were placed in two different locations with each trial in pseudo-random manner.

### Statistical analysis

We measured three main variables: total exploration time of each object, count of visits to each object and average exploration bout duration. Due to technical reasons, not all animals could be included in the count and bout duration analysis.

General behavioral differences during training were tested with a repeated measure ANOVA with day and session as factors for each variable separately. Additionally, we repeated the analysis but with condition instead of session (orthogonal thus not compatible in one analysis), which showed no significant effects. To test for memory for all three measure the discrimination index was used calculated as the difference in time exploring the novel object location and stable location divided by the total exploration time. This results in a score ranging from −1 (preference for the stable location) to +1 (preference for the less stable object location). A score of 0 indicates no preference for either object location. To test for the presence of memory, the discrimination index was test with one-sample t-test to chance. In case of non-normality of the data Wilcoxon Signed Rank Test was used.

### Model

The same computational model as in ^4^ was used (see article for more detailed methods).

In short, the model learns place-object associations and then translates this memory into an exploratory behavior: the objects that were stably found at the same location have a very low uncertainty and are thus less attractive during exploration than objects found at changing locations (high uncertainty in placeobject association).

The model employs two different parameters: a learning rate α, which determines the speed of memory accumulation; an inverse temperature β, which determines the strength and sign of memory expression during exploratory behavior.

A low learning rate α (i.e., close to 0) means that the model will need numerous repetitions of the same observation (i.e., in the Object Space Task, many trials observing the same place-object association) to properly memorize it. In contrast, a high learning rate α (i.e., close to 1) means that the model quickly memorizes new observations at the expense of old observations which more quickly forgotten. As a consequence, with a low learning rate the exploratory behavior generated by the model will mostly reflect remote memories but not recent ones (semantic-like memory). Conversely, with a high learning rate, exploratory behavior in the model will mostly reflect recent memories but not remote ones (episodic-like memory).

Finally, an inverse temperature β close to zero means that the model does not strongly translate memories into object preferences for exploration, thus showing little object preference. In contrast, a high inverse temperature will mean that the model’s exploratory behavior is strongly driven by differences in relative uncertainty between place-object associations. A high positive inverse temperature will result in neophilic behavior: the model spends more time exploring objects associated with high uncertainty (i.e., novelty or constantly changing location); a high negative inverse temperature will result in neophobic behavior: the model spends more time exploring objects with low uncertainty (stable/familiar objects). The model was fitted to each mouse’s trial-by-trial behavior using a maximum likelihood procedure described in [3]. In brief, this model fitting process found the best parameter values for each subject that best explain the relative proportion of time spent exploring each object at each trial. All model equations are described in [3].

### Automatic Behavioral Analysis

To classify genotype of mice in single trials, two main models were designed. The first model is a video action scoring classifier, upon which the general behavioural descriptions (i.e. features) were based. The second model is a genotype classifier that uses these general behaviours to predict WT/KO based on a single trial in the Object Space Task.

The model that scores mouse behaviours (Object Exploration, Wall Exploration, Corner Sitting) in a video is two-fold, in that one model classifies Object Exploration, whereas the other model classifies Wall Exploration and Corner Sitting. The first model is an inflated deep inception neural network based on Carreira and Zisserman ^31^ human kinetic action recognition network. Transfer learning was applied to a restructured version of the neural network and re-trained on videos of mice performing the Object Space Task, labelled by humans for object exploration. Next, the second model uses DeepLabCut ^18^ to extract limb configuration of mice over a single video. In addition to limb configuration, head direction was calculated as ears pointing to the nose. The limb configuration and head direction were then used in combination with knowledge about the square arena, such as location of walls and corners. Corner Sitting was defined as mean limbs location being near the corner, and Wall Exploration was defined as mean limb configuration being near a wall and head direction towards that wall.

The model that classifies genotype based on single trials, uses the behaviours extracted from the automatic behavioural scoring. To elucidate, behavioural summaries over the 5 minute trials were calculated based upon the time-series of actions (see Tab 1.). These features were then used as input variables for a Random Forest and XGBoost classifier, with genotype as a target variable. The classifiers were trained to optimize the AUROC (Area Under the Receiver Operating Curve), for each subset of data (all trials, stable trials, overlapping trials, random trials). Significance of the AUROC was then tested under the permutation distribution. To assess which behavioural features were driving most of each model in their predictions, feature importance were calculated was drop-column importance. The top features per classifier were taken for additional analysis, to further investigate how behaviours differ between WT and KO mice in single trials per condition. The mean difference was tested under the permutation distribution for each top feature, with genotype as a between-subject factor.

